# Habitat fragmentation induces rapid divergence of migratory and isolated sticklebacks

**DOI:** 10.1101/2021.08.20.457130

**Authors:** A. Ramesh, A.G.G. Groothuis, F.J. Weissing, M. Nicolaus

## Abstract

The adaptive capacity of many organisms is seriously challenged by human-imposed environmental change, which currently happens at unprecedented rates and magnitudes. For migratory fish, habitat fragmentation is a major challenge that can compromise their survival and reproduction. Therefore, it is important to study if fish populations can adapt to such modifications of their habitat. Here, we study whether originally anadromous three-spined stickleback populations (*Gasterosteus aculeatus;* ‘migrants’) changed in behavior and morphology in response to human-induced isolation. We made use of a natural field-experiment, where the construction of pumping stations and sluices in the 1970s unintendedly created replicates of land-locked stickleback populations (‘resident’) in the Netherlands. For two years, we systematically tested populations of residents and migrants for differences in morphology and behavioral traits (activity, aggressiveness, exploration, boldness and shoaling) in lab-based assays. We detected differences between migrant and resident populations in virtually all phenotypic traits studied: compared to the ancestral migrants, residents were smaller in size, had fewer and smaller plates and were significantly more active, aggressive, exploratory and bolder and shoaled less. Despite large ecological differences between 2018 and 2019, results were largely consistent across the two years. Our study shows that human-induced environmental change has led to the rapid and consistent morphological and behavioral divergence of stickleback populations in about 50 generations. Such changes may be adaptive but this remains to be tested.

**Lay summary:** The adaptive capacity of many organisms is seriously challenged by human-imposed environmental changes. For example, migratory fish encounter man-made barriers that impede their movements and force them to adopt a resident lifecycle. Here we study whether and how populations of three-spined sticklebacks diverged in response to human-induced isolation. We show that about 50 generations of isolation were sufficient to induce substantial morphological and behavioral differentiation between land-locked populations (‘residents’) and their migratory ancestors (‘migrants’).

## Introduction

Humans induce unprecedented fast changes in in many habitats, thereby imposing new selective pressures to animal populations (so-called ‘human-induced rapid environmental change’, sensu Sih, 2013). Animals thus need to implement quick adaptive responses to these changes to maintain their ability to survive and reproduce. One of the first responses to these challenges is often behavioral as behavior directly mediates how individuals interact with their environment. Consequently, it is expected that animal populations will respond to human-induced changes through behavioral modifications as a first step, which then may pave way for other morphological and/or physiological adaptations (Sih *et al*., 2011; Tuomainen & Candolin, 2011; Wong & Candolin, 2015).

Animal personalities are behavioral traits that are consistent across time or contexts and are often correlated to form “behavioral syndromes” (Stamps & Groothuis, 2010). Animal personalities presumably have significant consequences for the speed and the outcome of adaptation processes to changing environments (Bolnick *et al*., 2011; Dall *et al*., 2012; Sih *et al*., 2012; Wolf & Weissing, 2012). For example, personality variation may slow-down or speed-up rate of microevolution depending on whether personality structure retards adaptive evolution (Dochtermann & Dingemanse, 2013) or provides ‘pre-adapted’ phenotypes, which drive faster adaptation in multiple dimensions (Wagner & Altenberg, 1996; Barrett & Schluter, 2008; Wolf & Weissing, 2012; Van Gestel & Weissing, 2018). Furthermore, existence of personalities and mechanisms maintaining such intraspecific variation within populations can have an immense effect on the adaptive potential of these populations in response to environmental change (Réale *et al*., 2007; Bolnick *et al*., 2011; Dall *et al*., 2012; Wolf & Weissing, 2012; Moran *et al*., 2016).

Human-driven changes have disproportionately affected freshwater species, which have suffered the largest declines of 84% on average (WWF living planet report 2020). One of the greatest threats is habitat fragmentation that decreases habitat size and functional connectivity between habitats (Legrand *et al*., 2017). Migratory fish species, in particular, rely upon moving between sea and freshwater or between other habitats to reach spawning and nursery habitats (Fullerton *et al*., 2010). Hence, blocking access to these habitats can compromise the reproduction and survival of such migratory species (Hutchings, 2002). The important questions that connect the fields of animal personality, conservation, ecology, and evolution are whether and how migratory fish can adapt to the sudden isolation. Our study system in the north of the Netherlands is well suited to address such questions: In the last 50 years, man-made barriers (such as pumping stations and sluices) have been extensively built in rivers to maintain water levels below sea-level, with the consequence that it has blocked some of the side arms of main river channels. This created an unintended natural field experiment, wherein several populations of anadromous three-spined sticklebacks (*Gasterosteus aculeatus*) (‘migrants’) have become land-locked (‘residents’) in some of these replicate side-arms of the river. Over contemporary timescales, we expect resident populations of sticklebacks to have experienced very different selection pressures by completing their life-cycle entirely in freshwater as opposed to their ancestral migrants, that spend a significant part of juvenile growth at the sea, during winter. We used this opportunity to study whether resident populations exhibit consistent phenotypic differences (morphology and personality) compared to ancestral anadromous sticklebacks, as a result of this recent human-driven change.

Three-spined sticklebacks have become a model system for studying rapid phenotypic divergence because populations generally harbor high standing phenotypic variation (for instance, as reported in Jones *et al*. 2012), which enables them to adapt to a multitude of environments and through various proximate mechanisms (genetic, hormones, developmental plasticity, parental effects) (see review for freshwater colonization in Table 1). Likewise, it is repeatedly found that phenotypic differences can occur among populations with and without exposure to predation (Bell & Sih, 2007; Dingemanse *et al*., 2007, 2009; Stein *et al*., 2018; Dingemanse *et al*., 2020). Yet, little is known about population phenotypic divergence following habitat fragmentation over shorter timescales. To fill this knowledge gap, we sampled resident and migrant stickleback populations over two years and quantified differences in morphology and in behavioral traits involved in movements and anti-predator strategies: activity, aggressiveness, exploration, boldness and sociability (Magurran & Seghers, 1994; Cote *et al*., 2010, 2013; Chapman *et al*., 2011; Trompf & Brown, 2014; Sommer-Trembo *et al*., 2017, Wolf *et al*., 2008, 2011). In this species, these behaviors are moderately repeatable (for example, for the classical behavioral assays in Dingemanse *et al*., 2007, repeatability ranges between 0.339 – 0.552,) and can be phenotypically integrated (Bell & Stamps, 2004; Bell, 2005; Bell & Sih, 2007; Dingemanse *et al*., 2007, 2020; Kim & Velando, 2015). Our field system provides an good opportunity to answer (1) whether ∼50 years of isolation have been sufficient to induce morphological and behavioral differences between resident and migrant populations; (2) if the observed differences remained consistent over the two study years. Based on the synthesized literature on freshwater adaptation in this species (Table 1), we expect that individuals in resident populations should exhibit smaller body size with less armature as well as decreased levels of activity, exploration, boldness and shoaling compared to the ancestral migratory population.

**Table 1.**
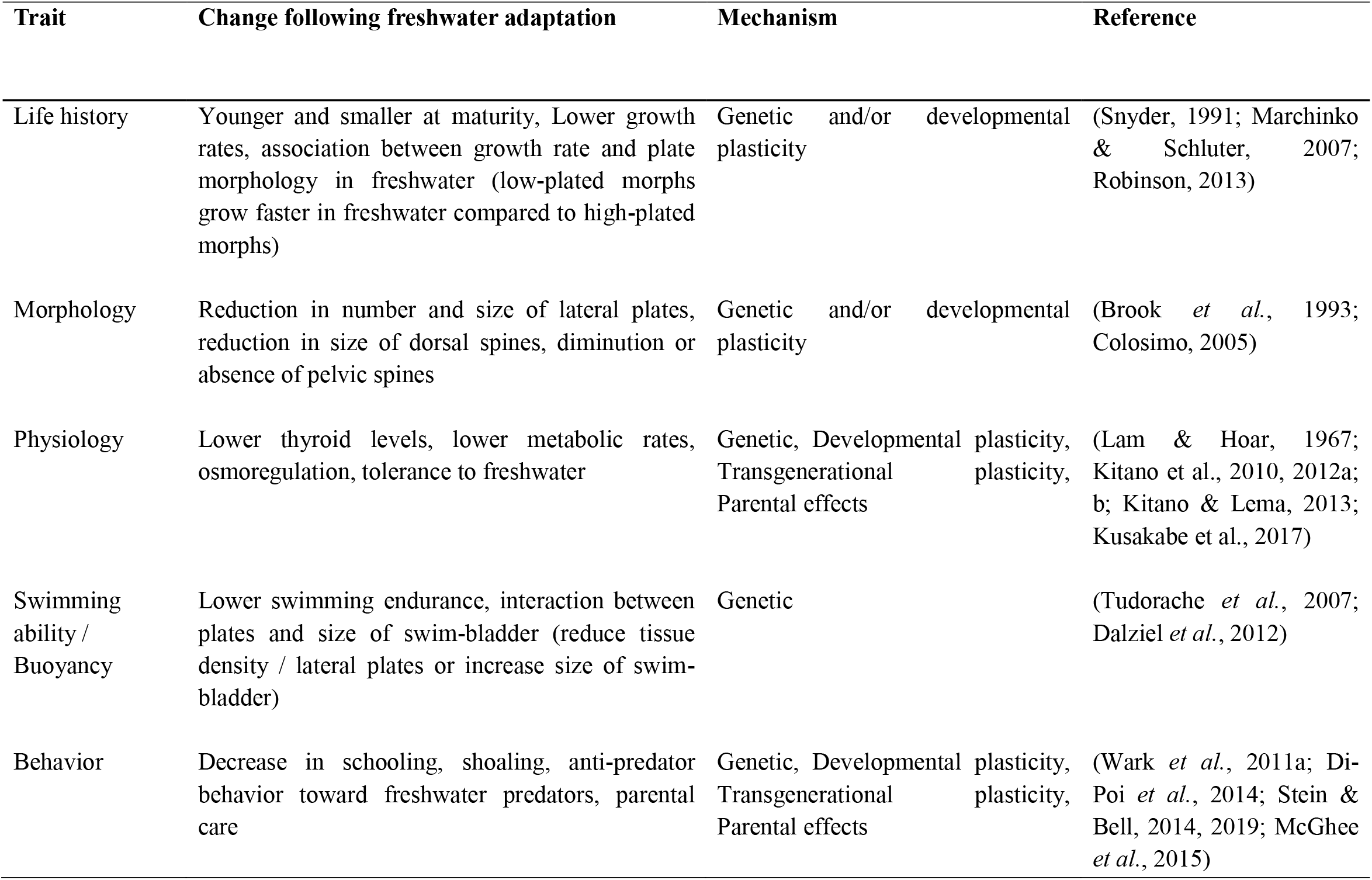
Table of freshwater adaptations from marine and migratory ancestors in three-spined sticklebacks.

## Methods

### Study populations and data collection

Our study sites were located along two main rivers, Termunterzijldiep and Westerwoldse Aa originating from the Ems Dollard estuary in the province of Groningen, the Netherlands. We caught incoming migrants at the two sea locks (“TER” (*53*^*0*^*18’7*.*24’’, 7*^*0*^*2’17*.*11’’*) and “NSTZ” (*53*^*0*^*13’54*.*49’’, 7*^*0*^*12’30*.*99’’*)), whereas resident sticklebacks were caught in two adjacent land-locked polders (“LL-A” (*53*^*0*^*17’56*.*14’’, 7*^*0*^*2’1*.*28’’*) and “LL-B” (*53*^*0*^*17’16*.*52’’, 7*^*0*^*2’26*.*46’’*)) (Supplementary information 1a). The land-locked populations are a part of a network of isolated freshwater ditches from the side arms of the main river, with depths less than three meters and width up to 8 meters with ample vegetation. LL-A is separated from the main river by a historic sluis (Supplementary information 1b) and LL-B is separated by a pumping station (Supplementary information 1c). The two land-locked locations are potentially connected but around 3 km apart in two locally different habitats; LL-A within wooded areas, with low human activity and LL-B in open farms with little shade and frequent human activity. To prevent sampling biases, we used lift-, hand- and fyke nets in resident populations and lift netting for incoming migrants directly at the fish passages at the sea locks, ensuring that fish of all behavioral types fish were caught. Apart from spatial separation, which aided us in assigning migrants and resident status to fish, we also we repeatedly sampled either in and outside the land-locked polders as well as in and outside of migratory season (with electrofishing) in order to confirm our assignment of ‘residents’. We consistently found resident fish all year round while migrants were absent outside the breeding season.

All individuals were transported to the laboratory within 2 hours of capture in aerated bags. After acclimatization to the laboratory conditions (temperature and 1 % salinity water) for one hour, we took the following morphological measurements of all individuals: total length (the length from the tip of the snout to the end of the tail), standard length (the length from the tip of the snout to the base of the tail), body mass, category of lateral plating (fully-plated, partially-plated and low-plated forms) (Bell & Foster, 1994) and clipped fins and/or spine of individuals for unique individual identification. We could not count the plates accurately and quickly in live specimens and used a rule of thumb that follows this description - low-plated (10 or fewer plates), partially-plated (11–20 plates), or completely-plated (21–30 plates). Low plates and fully plated forms were assigned with high accuracy, while semi-plated and fully plated ones were difficult to assign, especially in resident fish which are of considerable smaller size. We used *standard length* as proxy for size in all analyses because this measure is highly correlated with the two other measures namely, total length and body mass and was less error prone than total length (Supplementary information 3).

Thereafter, we placed each fish in an individual “home tank” (30 × 16 × 18 cm (L × W × H)) that was visually isolated from others and enriched with one artificial plant. Fish were fed frozen blood worms and brine shrimps (3F Frozen Fish Food bv.), *ad libitum*. On the following day, the fish were allowed to acclimatize to the new environment and laboratory conditions (day 0). From day 1 to day 4, fish were subjected to a range of behavioral tests (Fig. 1). On day 6 or 7 fish were released in the wild at their site of origin or kept in the lab for further breeding experiments. The laboratory conditions were set to mimic the natural conditions in terms of air temperature (range 5 to 20 ^0^C, depending on season) and photoperiod (range 10:14 L:D to 16:8 L:D, cycled with natural levels). Fish were monitored every day and individuals that showed signs of injury or sickness were removed immediately or before the behavioral assay (n=65).

**Figure 1.**
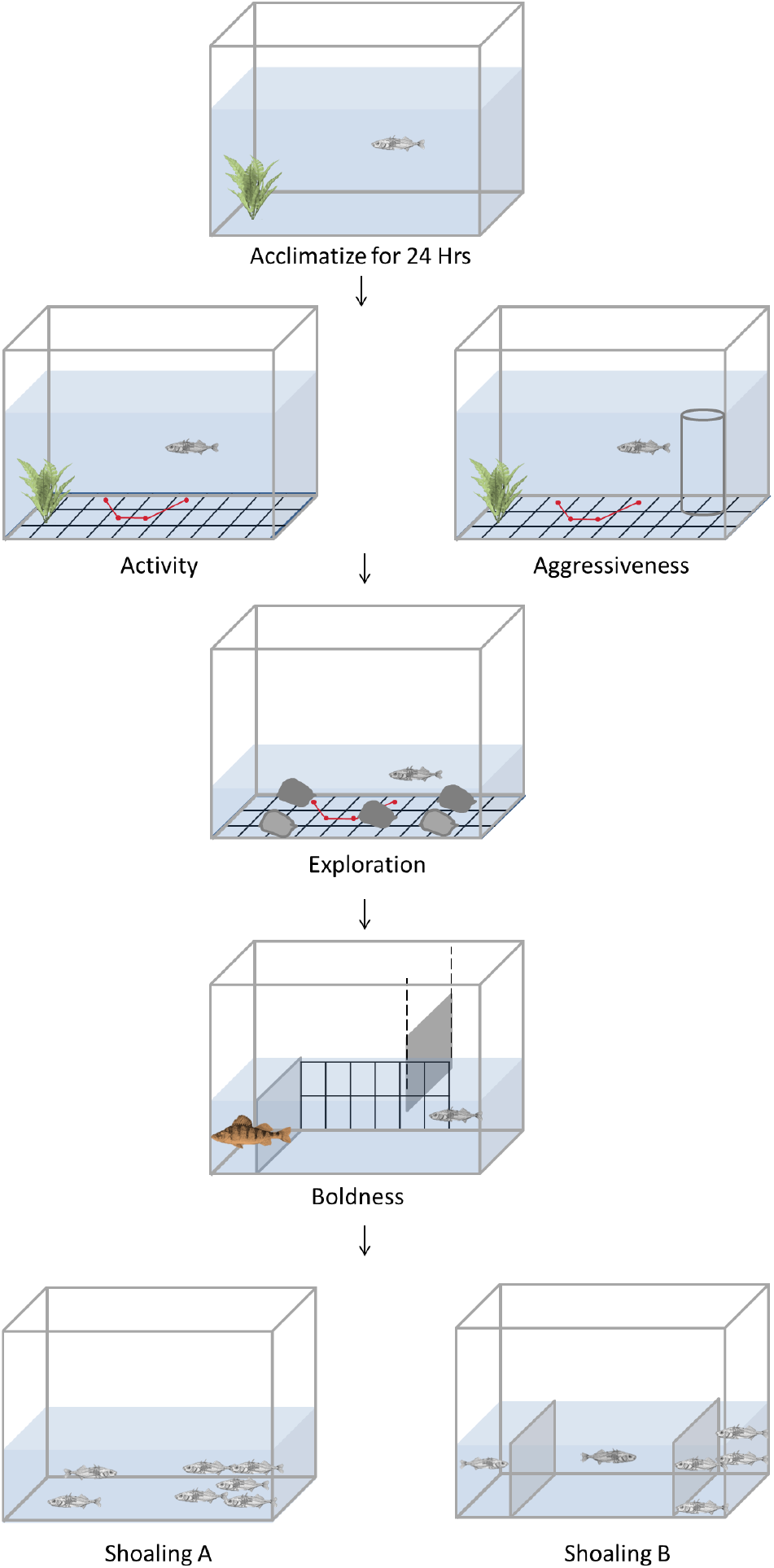
Behavioral assay. The flow chart represents the order in which assays were performed along with illustration of different behavioral assays and the placement of grids used for extracting different parameters.

Data collection occurred between March – May in the years 2018 and 2019. These years were drastically different in terms of the weather conditions of the summer and winter of the previous years (see Supplementary information 2). Compared to winter of 2017, the winter of 2018 was particularly cold with frozen ditches and main canals until March and the following summer was in contrast very warm and dry leading to small ditches partly drying up (Maximum daily temperature (2017 vs 2018): 29.9^0^C vs. 35.7^0^C ; and mean annual precipitation (2017 vs. 2018: 25.9cm vs. 17.88cm; data from Royal Netherlands Meteorological Institute). In 2018, a total of 251 fish were caught (189 migrants and 62 residents) and in 2019, 74 fish were caught (38 migrants and 36 residents). It is noted that in 2019, we were successful in capturing migrants from only one population (“NSTZ”). Our sample size was determined by the number of fish we could successfully catch, while ensuring that batches were caught at different time intervals to avoid confounding effects of season.

Wild animals were sampled using a fishing permit from *Rijksdienst voor Ondernemend Nederland* (The Netherlands) and an angling permit from the Hengelsportfederatie Groningen-Drenthe. Housing and testing of behaviors were in adherence to the project permit from the *Centrale Commissie Dierproeven* (The Netherlands) under license number *AVD1050020174084*.

### Behavioral assays

Five behaviors were scored for both migrants and residents: general activity in home tank, aggression towards a conspecific from the same population, exploration of a novel environment, boldness in a predator inspection trial and shoaling tendency (Fig.1). Activity, aggression and exploration were live-scored by five observers whereas the boldness tests and shoaling assays were filmed and subsequently scored using the software BORIS v.6.2.4. (Friard & Gamba, 2016). Details of each assay are given below. The tests were performed during the light period (usually between 9 am and 6 pm). The sequence of fish to be tested was drawn at random. It was not possible to be blind to the status of fish, as migrants and residents exhibited large size differences.

In the behavioral assays (except shoaling) and morphology, we measured several variables. We used one of these variables as a proxy of the behavior of interest, as it is easier to interpret. We thus performed principal component analyses (PCA) including all measured variables per behavior under study and used the first principal component PC1 (explaining most of the variance) as a proxy for the for activity and exploration (Supplementary information 3) and we chose other variables or a composite of it based on biological reasons for aggression and boldness, as mentioned below. The PCA-based results (not reported), did not differ from the results on the single variables.

### Activity

The general activity level of each individual was recorded in their home tank using a grid at the bottom of the tank (Fig. 1). Each individual was observed for a period of 60s and its position in a 10 × 6 square grid space was recorded every 5s. With the recorded position the following values were calculated: unique squares visited, number of square changes and total distance travelled (adapted from Dingemanse *et al*., 2007). In the analyses reported in the main text, we used *number of square changes* as a proxy for ‘activity’.

### Aggression

Immediately after the activity test, we introduced an empty transparent glass in one corner of the home tank, in order to acclimatize the focal fish to the new object (120s). Subsequently, the empty glass was replaced with a similar one containing a smaller conspecific from the same population (“intruder”). During the following 120s, we scored the position of the focal individual and its response towards the intruder (bites, spine-up display) every 10 seconds (Fig. 1). The mean and minimum distance to the intruder and the total number of bites were then calculated (adapted from Bell & Stamps, 2004). Spine-up threat display was hard to notice for residents because of their smaller spines and subsequently dropped from observations. We re-used intruders for a maximum of 5 different trials and controlled for intruder identity in the later analyses. To disentangle aggression from sociality, we used the *total number of bites* as a proxy for ‘aggression’, rather than the time spent near the intruder.

### Exploration

For studying exploration in a novel environment, the focal fish was placed into an opaque acclimatization compartment (4 × 6 cm) within a tank of size equal to the home tank, a water level of 5 cm, and with a 10 × 6 square grid at the bottom. The tank included five stones that extend to the top of the water surface to block the view and force the fish to swim around them to gather information about the environment (Fig. 1). After an acclimatization period of 120s, the compartment was gently removed, releasing the fish into the arena and the subject started the exploration test, lasting for 300s. During this period, the position of the focal fish was recorded every 5s. With the recorded position the following values were calculated: unique squares visited, number of square changes and total distance travelled (adapted from Dingemanse *et al*., 2007). In the analyses reported in the main text, we used *number of square changes* as a proxy for ‘exploration’.

### Boldness

In the boldness tests, we measured the responses of the focal fish toward a model of a predator, European perch (*Perca fluviatilis)*, with joined soft body that moves realistically when moved remotely using a thread (Kozak & Boughman, 2012). European perches naturally occur in our field sites and are considered one of the primary predators of sticklebacks (Hoogeland *et al*., 1956). The focal fish was moved from its home-tank into a bigger, novel tank (60 × 30 × 30 cm) with three compartments, filled with 10 cm of water. Of the three compartments, the predator model was presented in the left compartment while the focal fish was released from the right compartment. The space between the ‘fish’ compartment and ‘predator’ compartment was divided into 8 equally spaced grids with one fish-distance (6 cm) between the subsequent grids (Fig. 1). The focal fish was first placed into the fish compartment of the tank and the barrier at the fish compartment was removed while the opaque barrier at the predator compartment was retained. Subsequently, the focal fish could explore the novel tank for a period of 120s without the predator being visible. After that period, the focal fish was gently pushed back into the fish compartment and thebarrier was replaced. Meanwhile the opaque barrier to the predator model was removed and replaced by a transparent barrier. After this was done, the barrier of the fish compartment was removed again to allow focal fish to view predator and the boldness trial of 300s was recorded in a camera. In the subsequent video-scoring, the latency to exit the fish compartment, the number of inspection bouts (i.e. directed swimming towards the predator crossing at least one square and ending when the fish swam back into the opposite direction), the total duration of inspection bouts, the number of predator visits (i.e. visiting the last grid next to the predator compartment, <6 cm), the total duration spent near the predator compartment and the minimum distance to the predator compartment were recorded. If a fish did not exit, its latency to exit amounted to the maximum or 300s and all other values were recorded as NA (adapted from Wilson & Godin, 2009). At least half of the water was replaced after testing 10 fish in the arena.

We used the *number of inspection bouts* towards the predator (number of inspection bouts performed in the first minute after the focal fish entered the arena) as a proxy for ‘boldness’. This measure is preferable to *latency to exit*, as it is less related to activity, wherein a more active fish has higher probability to exit and to *time spent near predator*, as it takes into account the total time the fish spent in the test arena.

### Shoaling - A

In 2018, individual shoaling tendency was scored in a group of ten fish. Fish that were captured on the same day and within the same population were placed into a larger tank (60 × 30 × 30 cm) filled with 10cm of water where they could interact freely with each other (Fig. 1). After 120s, all shoaling fish and then all non-shoaling fish were caught, identified and shoal composition noted. Fish were considered to shoal if they associated with another fish within one-fish distance (< 6 cm) at the end of the test. The procedure was repeated three times to calculate a shoaling score or ratio (1.0 is when individual was found to be associated with the shoal in all three trials, adapted from Wark *et al*., 2011).

### Shoaling - B

The shoaling assay conducted in 2018 adopted a setting where individuals were able to interact with one another. However, this captured very little among-individuals differences. Hence, we readjusted this test in 2019 by assaying individual shoaling in a large tank divided into three compartments: a central testing arena where the focal fish was released and two end compartments containing the stimulus shoal (n=5 unfamiliar conspecifics) and two distracter fish (n=2 unfamiliar conspecifics) (Fig. 1; adapted from Wark *et al*., 2011). The stimulus shoal and distracter fish comprised of migrants if the focal fish was migrant and residents otherwise. The stimulus shoal and distracter fish compartments were switched in sides to prevent a place or side bias and the fish were replaced with new stimulus shoal and distractor fish after five trials. At the start of the test, the focal fish was allowed to acclimatize for 120s in the central arena without viewing the ends compartments that were covered with opaque barriers. The focal fish was returned to its home-tank momentarily and the opaque barriers were replaced with transparent barriers. The focal fish was then reintroduced to the center of the focal arena to record its shoaling behavior for 300s (shoaling time, spending one fish-distance (< 6 cm) from the shoal). The water was partially replaced after testing 10 fish in the arena.

### Statistical analyses

To test whether resident and migrant fish differ in the proportions of the three common lateral plate morphs, we used a Chi-squared test for each year separately. We then analyzed variation in standard length and all the behavioral traits measured (activity, aggression, exploration, standardized scores for predator inspection and shoaling) in Linear Mixed Models (LMMs) with Gaussian errors. *Status* (resident vs. migrant, with migrant being the reference category), *year* (2018 vs. 2019, 2019 being the reference category) and *status* **×** *year* interactions were included as fixed effects. Date and time of testing were not significant and were thus removed from the models. In all models, *observer ID* and the combination of *population ID-year* (four populations in 2018 and 3 populations in 2019, giving 7 levels, as population is the unit of replication for our inferences) were fitted as random effects. We did not detect any sex differences in any of the behaviors (Supplementary information 4), and thus decided to pool data from both sexes. All GLMMs were constructed in R v. 3.6.1, R Core Team (2019) using the lmer function of the ‘lme4’ package (Bates et al., 2015). The statistical significance of fixed effects was assessed based on the 95% confidence interval (CI): an effect was considered significant when its 95% CI did not include zero. The sample sizes slightly varied between tests due to missing data and are reported with the outcome of each statistical test.

To establish the existence and structure of behavioral syndromes in migrants and residents, we ran multivariate mixed models that estimate covariances and correlations among all traits. However, due to lack of model convergence, covariances could not be estimated this way. Other advocated methods (e.g. Structural equation modelling (SEM); Dingemanse *et al*., 2010) could not be applied due to limited sample sizes. Hence, we estimated syndromes based on pairwise Spearman correlation with sequential Benjamini and Hochberg correction for multiple testing (Benjamini & Hochberg, 1995). Data were zero-inflated in some behaviors (Aggression in residents; Activity and Aggression in migrants). We discarded these behaviors from the correlational analyses to prevent spuriously high correlation coefficients. Correlation analyses focus on pairwise relationships between traits, thus ignoring higher-order effects (Dingemanse *et al*., 2010). To overcome this, we also compared the results of a PCA approach to summarize the structure of relationship between all the behaviors within categories of migrants and residents between the years, which did not yield qualitatively different results (not shown).

## Results

### Morphological differentiation

Residents had more low-plated forms compared to migrants in 2018, but not in 2019 (Fig. 2a), although the difference between migrants and residents seem to display a similar pattern in both years. This is confirmed by chi-square test on the relative proportions of lateral plate morphs between residents and migrants (Proportions of fully, partial and low plated morphs in 2018 = 0.44, 0.47, 0.09 in migrants and 0.15, 0.30, 0.56 in residents respectively; χ^2^ (df=*2*, N = *247*) = *64*.*536*, p <0.01 and in 2019 = 0.68, 0.18, 0.15 in migrants and 0.56, 0.16, 0.28 in residents respectively; χ^2^ (df=*2*, N = *66*) = *1*.*785*, p = *0*.*410*). Residents were significantly smaller than migrants in both years (Table 1; Fig. 2b). All fish were also larger in 2019 compared to 2018 (Table 1; Fig. 2b).

**Figure 2.**
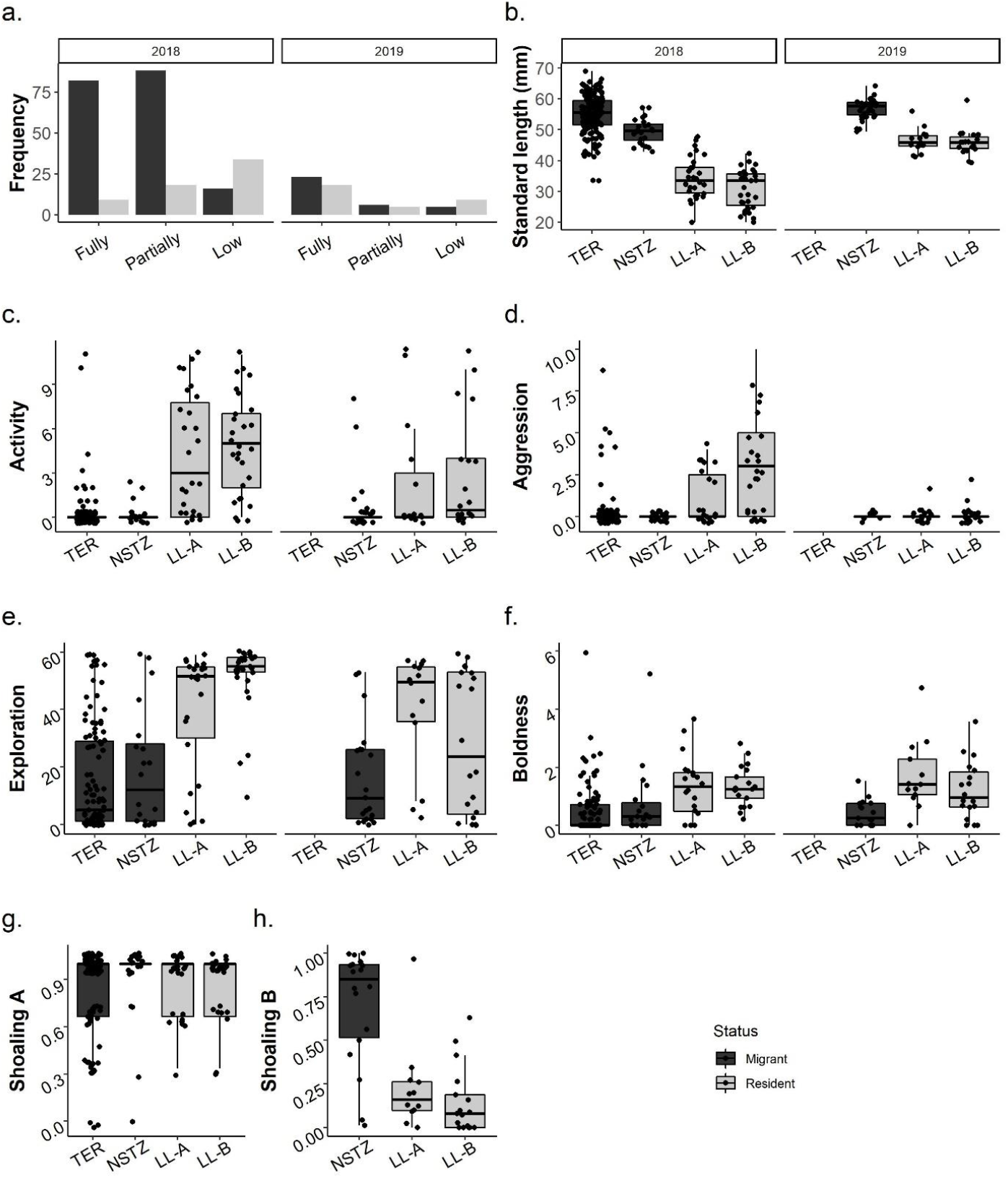
Individual size and behaviors (median ± quartiles) of two populations of residents and migrants over two years. **a**. Lateral plate morph distribution (N_2018_ =247, N_2019_= 66), **b**. Standard length ( N_2018_ =249, N_2019_= 72**), c**. Activity – number of square changes (N_2018_ =203, N_2019_= 56) **d**. Aggression – number of bites to intruder (N_2018_ =187, N_2019_= 44), **e**. Exploration – number of square changes (N_2018_ =183, N_2019_= 54), **f**. Boldness – number of inspection bouts / minute (N_2018_ =164, N_2019_= 48), **g**. Shoaling A – only 2018, fraction of trials spent with shoal (N_2018_ =180), **h**. Shoaling B – only 2019, fraction of time spent near stimulus shoal (N_2019_= 46)

### Behavioral differentiation

In both years, residents were significantly more active (87.5% of the migrants did not exhibit movements at all in their home-tanks), more exploratory and bolder compared to migrants (Table 2, Fig. 2c, e, f, respectively). Compared to previous studies in sticklebacks (Huntingford, 1976; Bell, 2005; Dingemanse *et al*., 2007), we found only a marginal proportion of aggressive individuals outside of the breeding period. In 2018, residents were significantly more aggressive than migrants and in 2019, this pattern disappeared (significant *status* **×** *year* in Table 2; Fig. 2d). The shoaling A assay performed in 2018 did not reveal differences between residents and migrants. However, the shoaling B assay performed in 2019 showed that residents shoaled much less than migrants (Table 2; Fig. 2h).

**Table 2.** Summary of linear mixed models on behaviors. For shoaling assays, the analyses were done separately for each year as they were different procedures and analyzed using linear models. Estimates of fixed effects (β) are given with their 95% confidence intervals (CI) computed by bootstrapping method and variance components are given with their standard deviation. Significant fixed effects are denoted in bold. Dashes represent random effects estimated at zero variance (models are singular). Other Sample size (N) represents number of individuals.

|  | Size<br>(N=317) | Activity<br>(N=258) | Aggression<br>(N=233) | Exploration<br>(N=239) | Boldness<br>(N=214) | Shoaling A<br>(N=181) | Shoaling B<br>(N=47) |
| --- | --- | --- | --- | --- | --- | --- | --- |
| *Fixed effects* | $\beta$ (95% CI) | $\beta$ (95% CI) | $\beta$ (95% CI) | $\beta$ (95% CI) | $\beta$ | $\beta$ | $\beta$ |
| Intercept | **52.428**<br>(48.551, 56.051) | 0.362<br>(-0.398, 1.108) | 0.181<br>(-1.258, 1.562) | **19.280**<br>(7.634, 30.471) | **0.754**<br>(0.080, 1.411) | **0.846**<br>(0.803, 890) | **0.706**<br>(0.580, 0.832) |
| Status (Resident) | **-19.808**<br>(-25.109, -14.505) | **4.337**<br>(3.402, 5.351) | **3.090**<br>(1.038, 5.239) | **29.014**<br>(16.430, 41.321) | **0.716**<br>(0.332, 1.064) | 0.036<br>(-0.043, 0.116) | **-0.526**<br>(-0.687, -0.366) |
| Year (2019) | 4.386<br>(-2.409, 11.190) | 0.614<br>(-0.534, 1.722) | 2.546<br>(-0.334, 5.555) | 2.466<br>(-16.140, 18.475) | **-0.166**<br>(-0.795, 0.475) |  |  |
| Status Resident x Year 2019 | **9.110**<br>(0.430, 17.688) | **-3.03**<br>(-4.799, -1.154) | **-5.367**<br>(-9.013, -1.501) | -17.183<br>(-37.133, 2.270) | **0.367**<br>(-0.509, 1.231) |  |  |
| *Random effects* | Variance (std.dev) | Variance (std.dev) | Variance (std.dev) | Variance (std.dev) | Variance (std.dev) |  |  |
| Observer ID |  | 0.171 (0.414) | - | 44.720 (6.687) | 0.350 (0.591) | - |  |
| Population - Year | 6.736 (2.595) | 0.000 (0.000) | 0.814(0.902) | 25.000(5.000) | - |  |  |
| Residuals | 32.780 (5.725) | 6.109 (2.472) | 10.138(3.184) | 346.810 (18.623) | 1.185 (0.920) | 0.062 (0.249) | 0.066 (0.256) |

### Behavioral syndromes

Behaviors were not correlated and there was little evidence for the existence of syndromes in both populations: only 2 of the 32 pair-wise correlations were significant after correcting for multiple testing and the correlation structure was not stable across years in either group (Fig. 3). In 2018, the only significant result was the positive correlation between exploration and predator inspection in migrants (Fig. 3, Supplementary information. 5; ρ = 0.29, corrected p = 0.009). In 2019, the only significant result was the positive correlation between activity and exploration in residents (Supplementary information. 5; ρ = 0.68, corrected p = 0.002). Most of the other correlations were far from significant.

**Figure 3.**
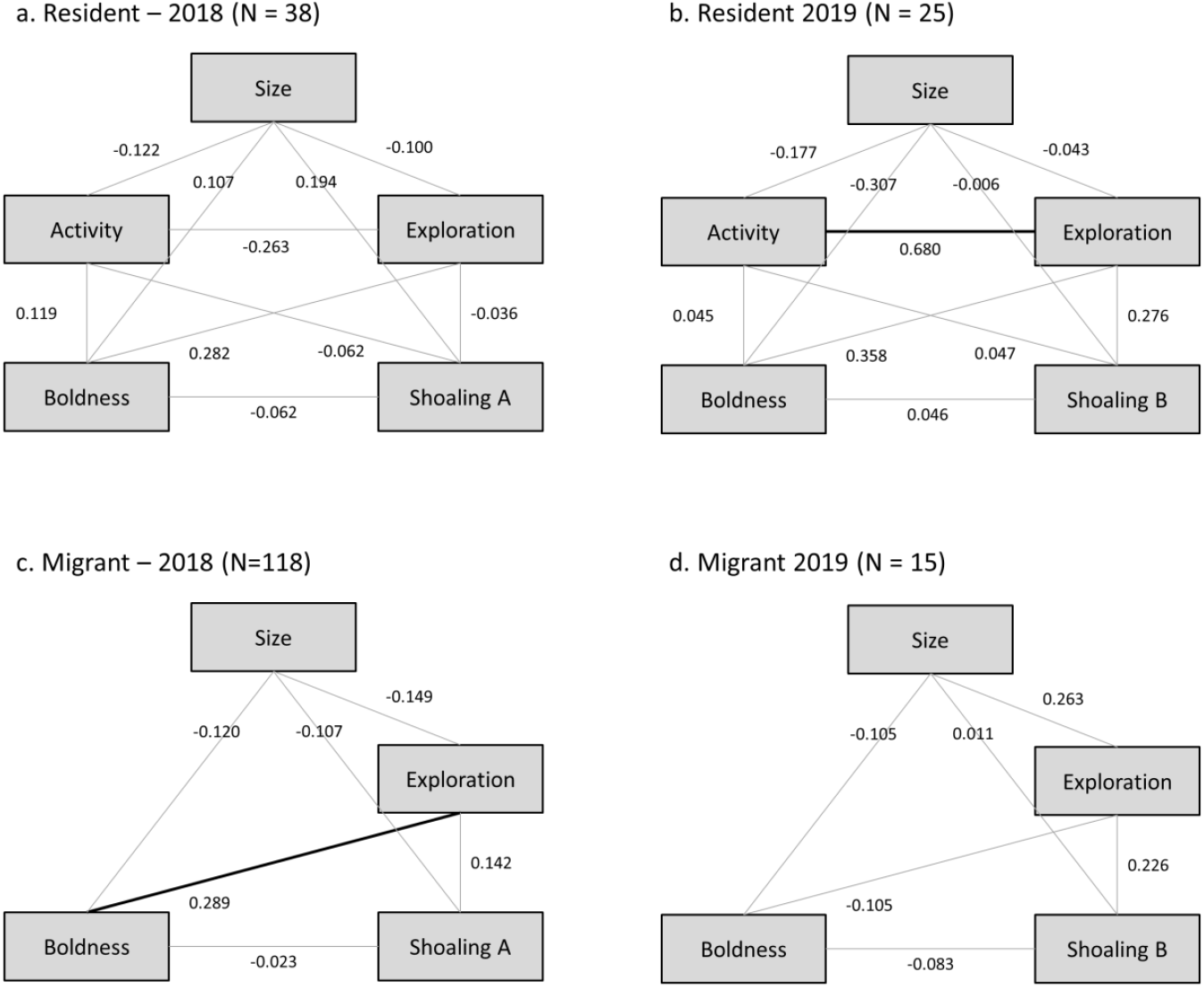
Syndrome structure of migrants and residents in two years. Significant correlations after sequential Benjamini Hochberg correction are represented with bold black lines. The numerical values represents pairwise Spearman correlation coefficients (rho).

## Discussion

This study investigated if resident populations of sticklebacks, which are cut off from the sea due to human water management measures in the 1970s, exhibit consistent morphological and behavioral differences compared to their ancestral migrant counterparts. Our results reveal that ∼50 generations of isolation were sufficient to induce substantial morphological and behavioral differences.

### Phenotypic divergence between derived residents and ancestral migrants

We found clear phenotypic differentiation between migrants and residents in almost all traits studied in both years. In line with previous literature on morphological adaptations of sticklebacks to freshwater that occurred over the last glaciation event (∼12,000 years), we found that residents were about half the size of migrants and were characterized mostly by low-plated forms. Although the resident fish from 2019 comprised of more fully plated forms than the residents in 2018, they showed reduction in plate width (personal obs.) in contrast to the robust armature spanning the width of the body, in fully- or partially-plated migrants. Full and shorter plates have also been observed in other freshwater populations, for example in freshwater lakes in Haida gwaii, low to fully-plated morphs are present depending on the presence of predators and turbidity of water (Reimchen *et al*., 2013) and depending on differential predator regimes and openness of habitats in the Baltic Sea (Eriksson *et al*., 2021). In our case, it suggests another pathway of reduction in lateral plate coverage by reducing the width of plates, as observed in populations in which the allelic variation for low-plated morph is limiting (Leinonen *et al*., 2012).

The morphological difference between populations are most easily explained by the necessity for flexibility to maneuver through vegetation in residents as compared to the demanding robustness and swimming abilities for migrants (Tudorache *et al*., 2007; Dalziel & Schulte, 2012; Dalziel *et al*., 2012) and decreased resource availability in freshwater during growth (Wund *et al*., 2012; Snyder, 1991). Furthermore, the reduction in number and size of lateral plates are also known to occur in response to different predator regimes present in the freshwater system (with fewer piscivorous predators and mainly dominated by invertebrate predators like dragonfly naiads) through selection on *eda* gene underlying lateral plate polymorphism (Colosimo *et al*., 2005; Marchinko & Schluter, 2007; Leinonen *et al*., 2011). These observations therefore suggest that the reduced size and the reduced armament of our resident fish likely follow the same pattern of adaptive evolution seen in other marine-freshwater stickleback systems.

As for individual behavioral scores, we found that residents were more active, aggressive (in the year 2018), exploratory, bolder and showed lower shoaling tendencies than migrants (in the year 2019). The majority of our findings with wild-caught sticklebacks are in line with the only other study that compared similar behaviors in populations of residents and migrants in lab-bred F1 sticklebacks (Di-Poi *et al*., 2014). In this study, the authors found that residents were more active, more aggressive and shoal less than migrants. Functional explanations for the behavioral differences can be given, but they include quite some speculation. Compared to the sea, land-locked ditches in our study sites are characterized by small and shallow streams, enriched with vegetation, low mean annual productivity (Gross *et al*., 1988), lower density of piscivorous fish yet with the presence of invertebrate predators (Reimchen, 1980; Marchinko, 2009) and birds. Hence for residents selection may favor higher levels of aggression and exploration that facilitate the discovery, acquisition and monopolization of limited resources (Budaev, 1997; Brown *et al*., 2005; Huizinga *et al*., 2009; Herczeg *et al*., 2013; Greenwood *et al*., 2016; Moran *et al*., 2017). Such ‘risk-prone’ behaviors may then be traded-off against shoaling, explaining why residents shoaled less compared to migrants (Ward *et al*., 2004). Differences in shoaling tendencies may also stem from the fact that migratory lifestyle involves group schooling during migration and presumably high shoaling tendencies in the sea due to ‘openness’ of habitats. In migrants, lowered activity level could further be an indication that freezing is an adaptive response to higher perceived predation when not protected by a shoal. (Huntingford & Wright, 1993). Furthermore, the robust armature and larger spines, characteristic of migrants, are known to impede them in escape behavior, thus potentially favoring freezing behavior (Andraso & Barron, 1995). In addition, reduced aggressive interactions could be due to the highly shoaling lifestyle of migrants as these two behaviors were shown to be incompatible in sticklebacks (Lacasse & Aubin-Horth, 2014). Despite the substantial differences in ecological conditions across the two study years, the differences in morphology and behavior between migrants and residents were relatively consistent, suggesting that the observed population differences are related to the different life styles of migrants and residents. We are aware that these results are obtained for stickleback populations that have been isolated for at least 50 generations but could be potentially diverging for longer. For example, in our migrant-resident stickleback system, divergence may have started prior to the construction of pumping station if behavioural polymorphism leading to partial migration is present within populations (McKinnon et al., 2004). Furthermore, the two resident populations in our study are separated from the migrants by a pumping station and a sluice (Supplementary information 1b,c). Although pumping stations pose as impenetrable barriers to adult stickleback, juveniles and fry may potentially cross over, and consequently reduced gene flow is possible. There exist many other possibilities and it is still remarkable that the stark behavioural differences in wild-caught fish from these resident and migrant populations exist, despite the potential of reduced gene flow to hamper local adaptation (Raeymaekers et al., 2014). To confirm this, genetic studies would be needed to quantify population divergence.

### Rates of phenotypic change in contemporary timescales

Though we find substantial phenotypic differentiation, the question still remains if the rates of differentiation are fast and approaching contemporary evolution. The current literature on the impact of anthropogenic changes leading to population differentiation (Hendry *et al*., 2008) and specifically in sticklebacks acts as a useful yardstick in this respect, however mostly in morphological traits. A useful way to quantify the rates of change is using haldanes (Haldane, 1949; Gingerich, 1993). Two classical ways of calculating this is using percentage change in trait means per generation and absolute change in trait standard deviations per generation. In our study, we found that rate of change in size was -0.007 haldanes or size decreased by 0.007 SDs upon freshwater isolation of migrants which is much smaller than what is reported in another anadromous-freshwater system (0.234 haldanes for females and 0.365 haldanes for males, Baker *et al*., 2011). Furthermore, the rates of changes in other behaviors upon isolation were 0.01 haldanes for activity, 0.001 haldanes for exploration, 0.014 haldanes for boldness and -0.149 haldanes for shoaling behavior. These rates compare to the rates of contemporary evolution of other traits in sticklebacks (ranging from 0.01 to 0.473 haldanes, Bell & Aguirre, 2013) and other organisms (0.0 to 1.142 haldanes, Hendry et al., 2008). In our studies, we see that rates of change of most behaviors are higher than that for size, which is as expected for traits. However, we should note overall that much of the differentiation we study are behavioral and can be brought about in the first few generations after isolation and our rates of change are underestimated as they are calculated linearly over 50 years. Indeed more recent studies on sticklebacks isolated from marine to freshwater habitats have found evidence for evolution on contemporary timescales of decades to even seasons (Lescak *et al*., 2015; Hosoki *et al*., 2019; Garcia-Elfring *et al*., 2021). It is to be noted that our values on rates of change are still preliminary because we do not know if these changes reflect genetic differentiation.

### Population differences in syndromes

A previous study with twelve freshwater stickleback populations reported a positive correlation between boldness and aggression toward a conspecific in five out of the six populations where predators were present (Dingemanse *et al*., 2007, 2009). There were also tight correlations among other behaviors including activity, exploration, aggressiveness and boldness in predator-sympatric populations (Correlation coefficients range from 0.03 to 0.74). These tight behavioral correlations are thought to result from predation that enhances habitat heterogeneity by creating risky and non-risky areas and thus favors alternative behavioral strategies (for example, Bell & Sih, 2007; Dingemanse *et al*., 2007; Dhellemmes *et al*., 2020). Surprisingly, (but in line with an earlier study on freshwater and marine sticklebacks, Di-Poi et al 2014), none of our stickleback populations, including migrants that should be exposed to higher predation pressure, exhibited stable syndromes across years and only few correlations between traits were detected. Boldness-Exploration was one of the stronger correlations in migrants (ρ = 0.289), but still was weaker compared to previous studies (ρ = 0.667, Dingemanse et *al*., 2007). Activity – Exploration syndrome in residents was observed in the second study year (ρ = 0.680), which was comparable to those reported from predator-sympatric populations (ρ = 0.754, Dingemanse et *al*., 2007). This lack of syndromes could be because the behaviors selected are not under correlated selection or that we lack the power to detect syndromes. Alternatively, in our system, predation risk and change in life-history may not systematically select for phenotypic trait integration (Sommer-Trembo *et al*., 2017).

## Conclusions

We have shown that behavior and morphology diverged in sticklebacks after human disturbance, blocking migration over about 50 generations. The observed phenotypic differences between migrants and residents clearly show that barriers to migration have thus major consequences for the phenotype and potentially life-histories and population dynamics of sticklebacks as correlated life-history characteristics (growth rate, size at maturity, number and size of eggs) are also known to change on adaptation to freshwater in sticklebacks. Nevertheless, at least some populations can cope to a drastic loss of migration opportunity as they seem to thrive in land locked conditions. Next step would be to test whether the observed divergence is adaptive and to identify how it came about. One way to delineate the relative roles of genetic inheritance, non-genetic inheritance, developmental plasticity and phenotypic plasticity is through common-garden experiments combined with cross-fostering experiments and through experiments where juveniles are exposed to different selective regimes in semi-natural mesocosms. This would give us insight into role of personality in adaptation to novel environmental conditions.

## Acknowledgements

We thank Louis de Vries, Colleen Illing, Diederik Blaauw, Lisette Borchers for helping with data collection, the animal care takers and Suma Murthy for helping with animal care and lab work. We thank Peter Paul Schollema, at the Water Authorities Hunze en Aa’s and Jeroen Huisman at van Hall Larenstein, university of applied sciences for help with acquiring samples of sticklebacks. We thank Jakob Gismann for commenting on the manuscript. We also thank two anonymous reviewers for their useful comments, which greatly improved the manuscript.

## Supplementary information of the manuscript entitled

**Supplementary information 1:**
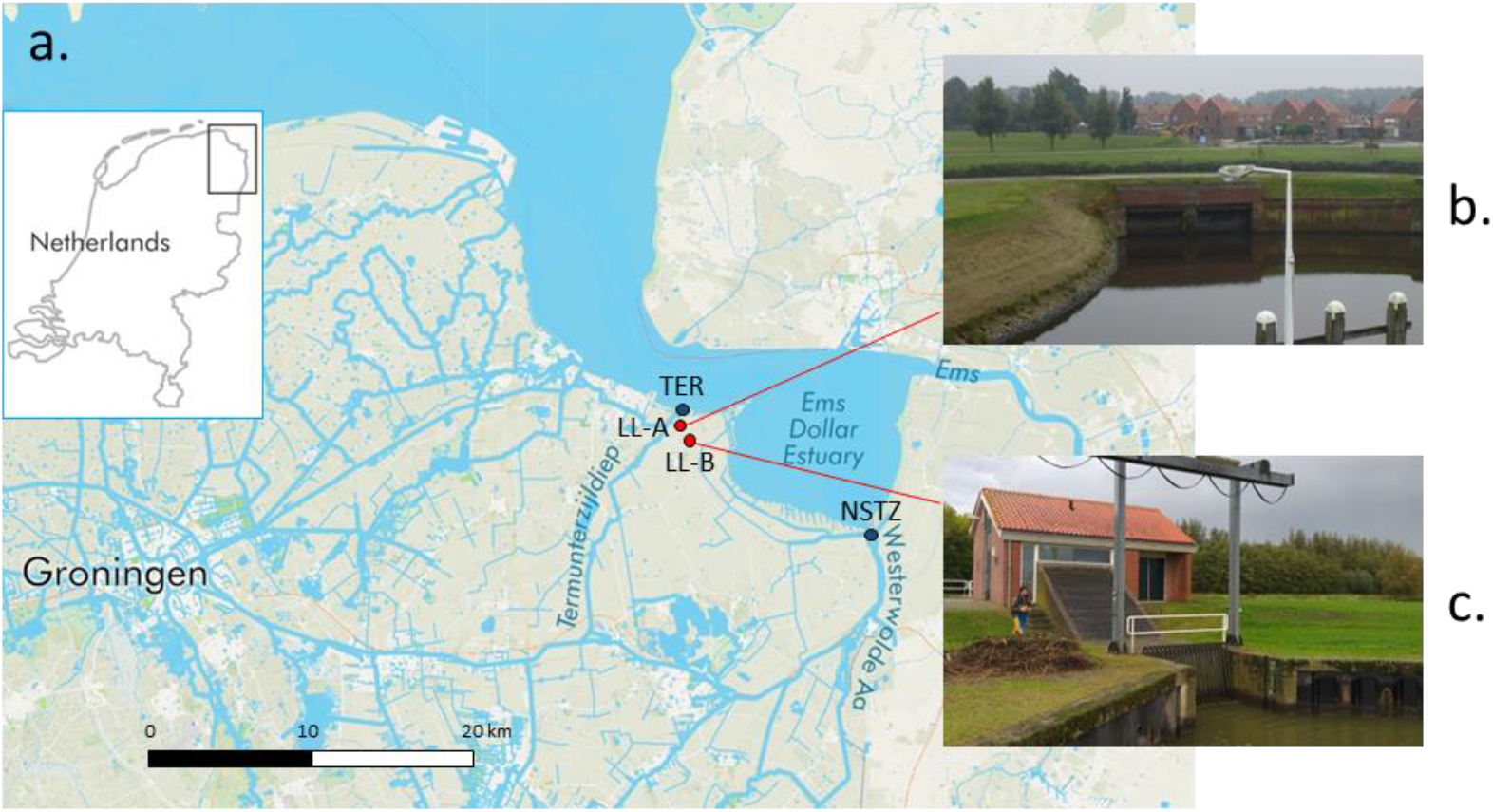
Map of the sampling sites and images of the barriers to migration in front of land-locked sites. a). Blue dots indicate the two open sites TER and NSTZ where incoming migrants were caught and red dots indicate the two land-locked sites LL-A and LL-B. b) LL-A is blocked by a historic sluis, which has not been operational for the last 50 years. c). LL-B is blocked by a pumping station, which allows water flow but restricts fish movement. Photo courtesy: P.P. Schollema, Waterschap Hunze en Aa’s

**Supplementary information 2:**
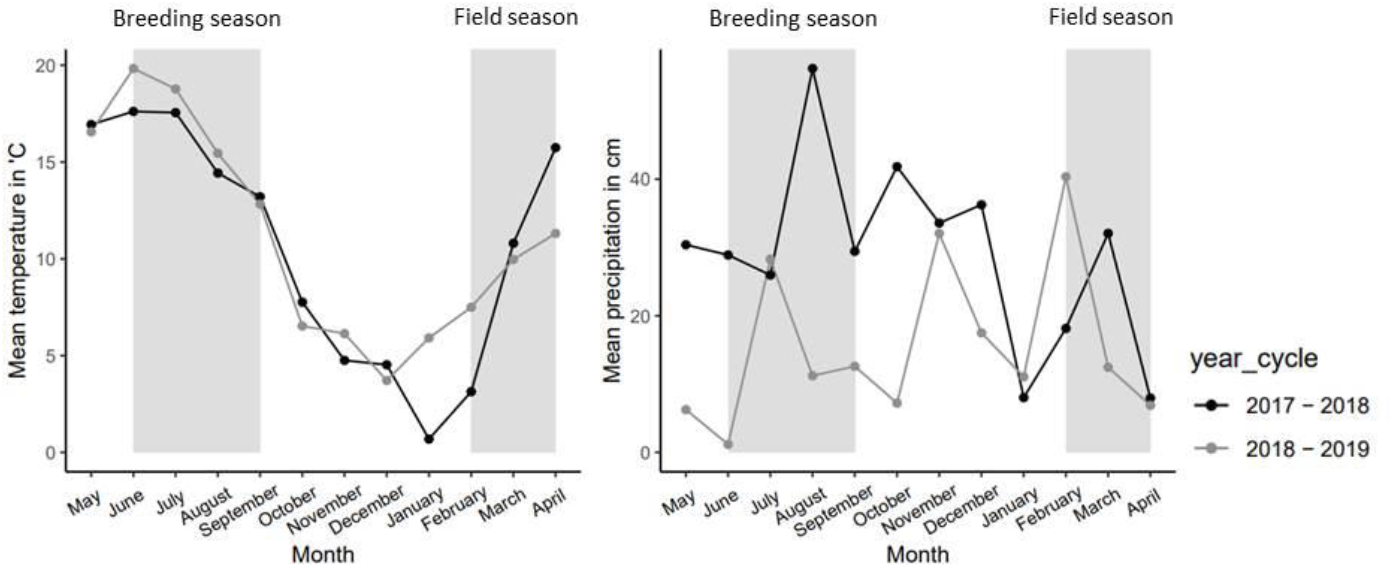
**Mean monthly temperature and precipitation of two year-cycles of weather station Lauwersoog (53. 02’37.6’’, 60. 12’44.8’’), which closely represents the weather conditions of the coastal areas in the north of the Netherlands**. Field season refers to the migratory period during which sticklebacks are caught. Breeding season refers to the breeding window, which coincides with spring and summer.

**Supplementary information 3:**
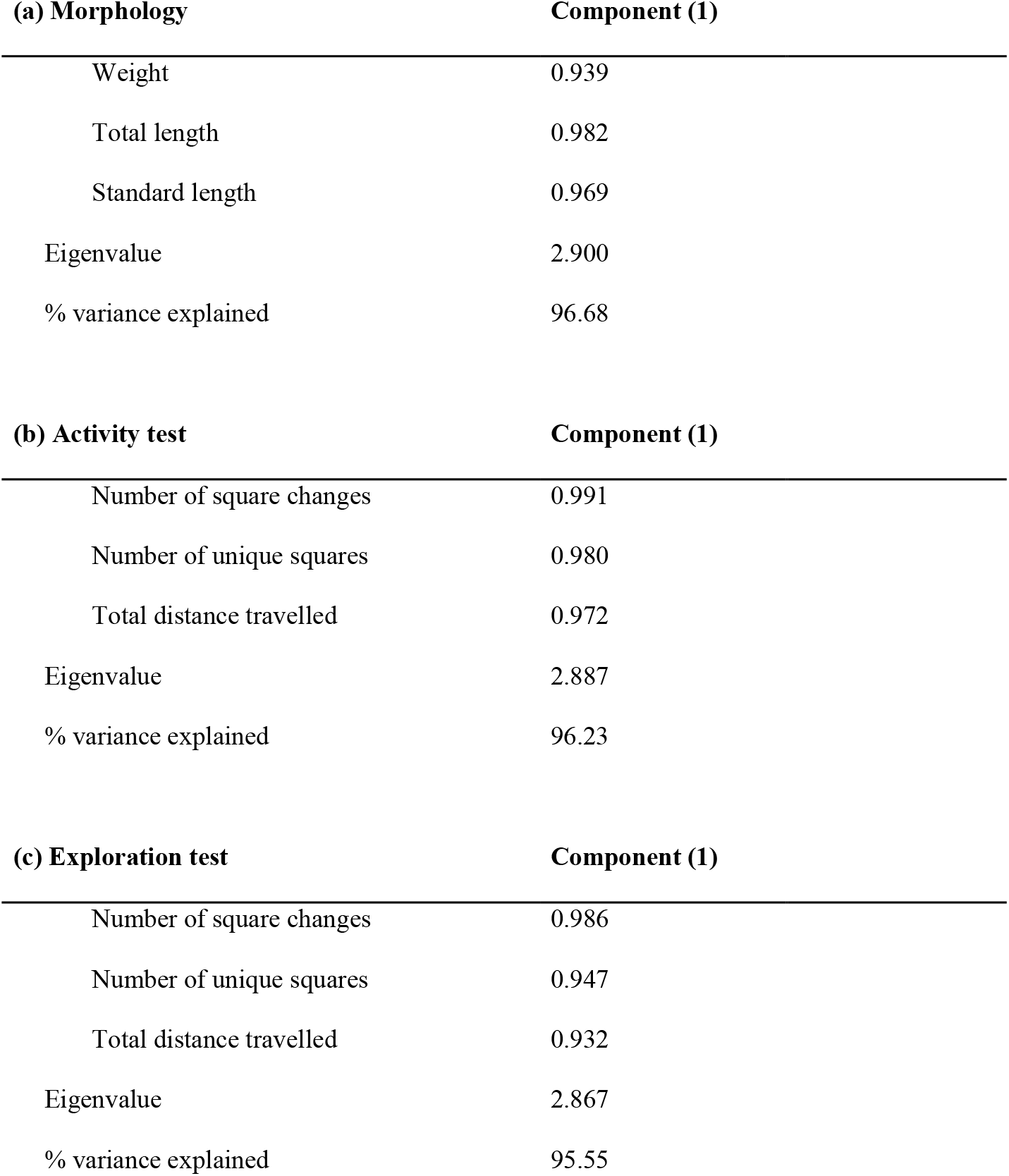
Results of principal component analysis on morphology and individual behaviors containing more than one variable: The trait, its variables and their corresponding cos2 value (or) contribution to the principal components along with the percentage of variance explained by each principal component are given.

| **(a) Morphology** | **Component (1)** |
| --- | --- |
| Weight | 0.939 |
| Total length | 0.982 |
| Standard length | 0.969 |
| Eigenvalue | 2.900 |
| % variance explained | 96.68 |

| **(b) Activity test** | **Component (1)** |
| --- | --- |
| Number of square changes | 0.991 |
| Number of unique squares | 0.980 |
| Total distance travelled | 0.972 |
| Eigenvalue | 2.887 |
| % variance explained | 96.23 |

| **(c) Exploration test** | **Component (1)** |
| --- | --- |
| Number of square changes | 0.986 |
| Number of unique squares | 0.947 |
| Total distance travelled | 0.932 |
| Eigenvalue | 2.867 |
| % variance explained | 95.55 |

**Supplementary information 4:**
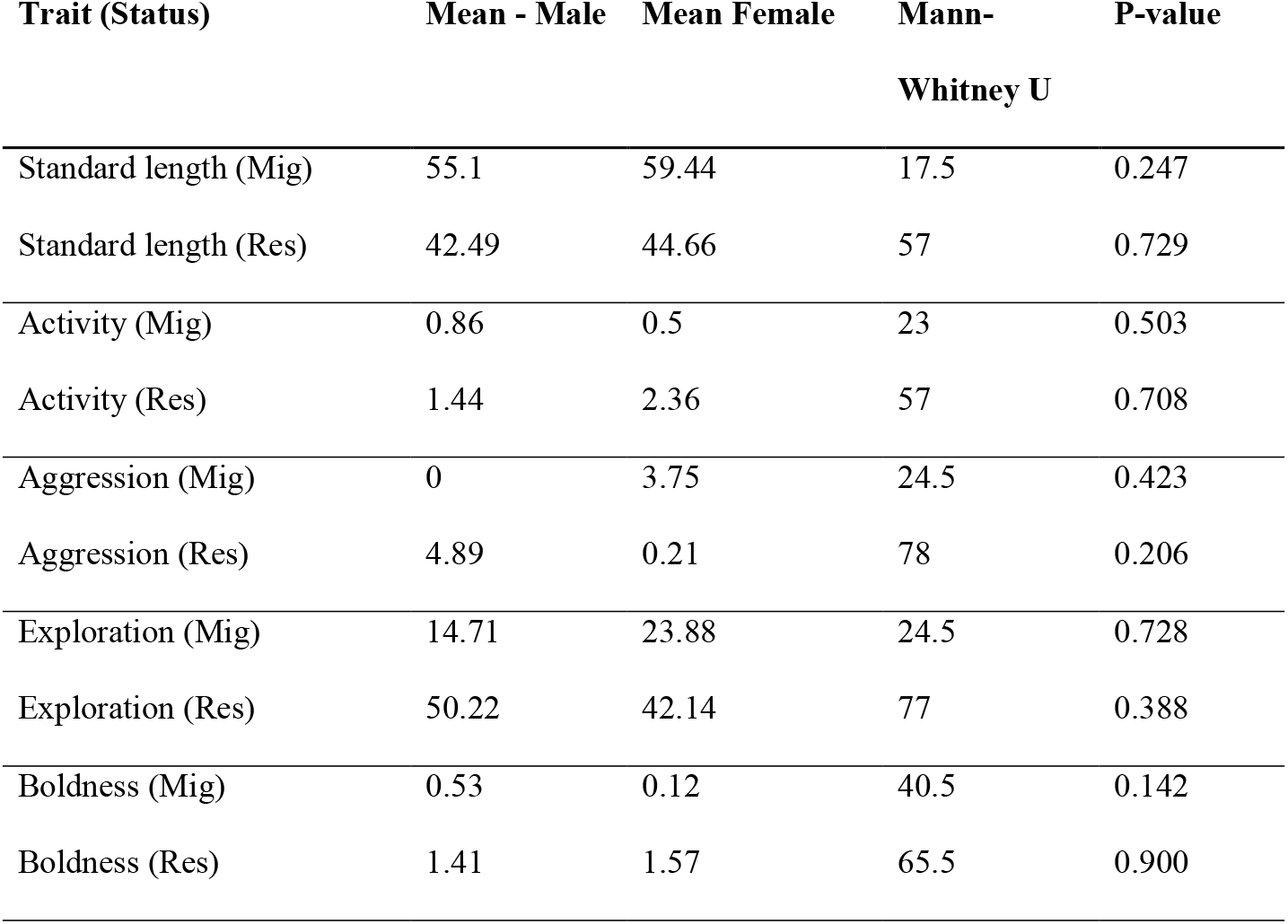
Behavior does not significantly differ between the sexes. Means and corresponding Mann-Whitney U statistic for size and individual behaviours, are given separated for migrants and residents females (N_migrant_=8 females and 7 males, N_resident_=14 females and 8 males). Shoaling was not compared as we had limited sample size for the two different shoaling assays.

**Supplementary information 5:** Correlations between behaviors and body size per status and year. The different behavior-behavior and behavior – morphology pairs are correlated using Spearman’s correlation coefficient along with Benjamini and Hochberg correction for multiple testing. Significant correlations after correction are denoted in bold.

| Resident 2018 (N=38) |  |  |  | Migrant 2018 (N=118) |  |  |  |
| --- | --- | --- | --- | --- | --- | --- | --- |
| Correlations | Spearman’s | P-Value | Corrected | Correlations | Spearman’s | P-Value | Corrected |
|  | rho |  | P-value |  | rho |  | P-value |
| Act – Exp | 0.263 | 0.111 | 0.555 | **Exp – Pred** | **0.289** | **0.0015** | **0.009** |
| Act – Pred | 0.119 | 0.475 | 0.786 | Exp – Shoal | 0.142 | 0.124 | 0.248 |
| Act – Shoal | -0.062 | 0.712 | 0.791 | Exp - Size | -0.149 | 0.106 | 0.248 |
| Act – Size | -0.122 | 0.464 | 0.786 | Pred – Shoal | 0.023 | 0.803 | 0.803 |
| Exp – Pred | 0.282 | 0.087 | 0.555 | Pred – Size | -0.120 | 0.195 | 0.293 |
| Exp – Shoal | -0.036 | 0.830 | 0.83 | Shoal - Size | -0.107 | 0.247 | 0.296 |
| Exp - Size | 0.100 | 0.550 | 0.786 |  |  |  |  |
| Pred – Shoal | -0.062 | 0.711 | 0.791 |  |  |  |  |
| Pred – Size | 0.107 | 0.522 | 0.786 |  |  |  |  |
| Shoal - Size | 0.194 | 0.243 | 0.786 |  |  |  |  |

| Resident 2019 (N=25) |  |  |  | Migrant 2019 (N=15) |  |  |  |
| --- | --- | --- | --- | --- | --- | --- | --- |
| Correlations | Spearman’s | P-Value | Corrected | Correlations | Spearman’s | P-Value | Corrected |
|  | rho |  | P-value |  | rho |  | P-value |
| **Act –Exp** | **0.680** | **0.0002** | **0.002** | Exp – Pred | -0.105 | 0.71 | 0.923 |
| Act – Pred | 0.045 | 0.831 | 0.930 | Exp – Shoal | 0.226 | 0.417 | 0.923 |
| Act – Shoal | 0.047 | 0.824 | 0.930 | Exp - Size | 0.263 | 0.343 | 0.923 |
| Act – Size | -0.177 | 0.398 | 0.796 | Pred – Shoal | -0.083 | 0.769 | 0.923 |
| Exp – Pred | 0.358 | 0.079 | 0.395 | Pred – Size | -0.150 | 0.594 | 0.923 |
| Exp – Shoal | 0.276 | 0.182 | 0.455 | Shoal - Size | 0.011 | 0.970 | 0.970 |
| Exp - Size | -0.043 | 0.837 | 0.930 |  |  |  |  |
| Pred – Shoal | 0.046 | 0.827 | 0.930 |  |  |  |  |
| Pred – Size | -0.307 | 0.136 | 0.453 |  |  |  |  |
| Shoal - Size | -0.006 | 0.977 | 0.977 |  |  |  |  |

## References

Andraso, G.M. & Barron, J.N. 1995. Evidence for trade-off between defensive morphology and startle-response performance in the brook stickleback (Culaea inconstans). Can. J. Zool. 73: 1147–1153.

Baker, J.A., Heins, D.C., King, R.W. and Foster, S.A. 2011. Rapid shifts in multiple life history traits in a population of threespine stickleback. J. Evol. Biol. 24: 863–870.

Bell, A.M. & Sih, A. 2007. Exposure to predation generates personality in threespined sticklebacks (Gasterosteus aculeatus). Ecol. Lett. 10: 828–834.

Bell, A.M. 2005. Behavioural differences between individuals and two populations of stickleback (Gasterosteus aculeatus). J. Evol. Biol. 18: 464–473

Bell, A.M. & Stamps, J. 2004. Development of behavioural differences between individuals and populations of sticklebacks, Gasterosteus aculeatus. Anim. Behav. 6: 1339–1348

Bell, M.A. & Aguirre, W.E. 2013. Contemporary evolution, allelic recycling, and adaptive radiation of the threespine stickleback. Evol. Ecol. Res. 15: 377–411.

Bell, M.A. & Foster, S.A. 1994. The evolutionary biology of the threespine stickleback. Oxford University Press.

Bell, M.A., Ortí, G., Walker, J.A., Koenings, J.P. 1993. Evolution of pelvic reduction in threespine stickleback fish:A test of competing hypotheses. Evolution. 47: 906–914.

Bolnick, D.I., Amarasekare, P., Araújo, M.S., Bürger, R., Levine, J.M., Novak, M., et al. 2011. Why intraspecific trait variation matters in community ecology. Trends Ecol. Evol. 26: 183–192.

Brown, C., Jones, F. & Braithwaite, V. 2005. In situ examination of boldness-shyness traits in the tropical poeciliid, Brachyraphis episcopi. Anim. Behav. 70: 1003–1009.

Budaev, S. V. 1997. Alternative styles in the European wrasse, Symphodus ocellatus: Boldness-related schooling tendency. Environ. Biol. Fishes 49: 71–78.

Colosimo, P.F., Hosemann, K.E., Balabhadra, S., Villarreal, G., Dickson, H., Grimwood, J., et al. 2005. Widespread parallel evolution in sticklebacks by repeated fixation of ectodysplasin alleles. Science. 307: 1928–1933.

Dall, S.R.X., Bell, A.M., Bolnick, D.I. & Ratnieks, F.L.W.W. 2012. An evolutionary ecology of individual differences. Ecol. Lett. 15: 1189–1198.

Dalziel, A.C. & Schulte, P.M. 2012. Correlates of prolonged swimming performance in F2 hybrids of migratory and non-migratory threespine stickleback. J. Exp. Biol. 215: 3587–3596.

Dalziel, A.C., Vines, T.H. & Schulte, P.M. 2012. Reductions in prolonged swimming capacity following freshwater colonization in multiple threespine stickleback populations. Evolution. 66: 1226–1239.

Dhellemmes, F., Finger, J.S., Laskowski, K.L., Guttridge, T.L. & Krause, J. 2020. Comparing behavioural syndromes across time and ecological conditions in a free-ranging predator. Anim. Behav. 162: 23–33.

Di-Poi, C., Lacasse, J., Rogers, S.M. & Aubin-Horth, N. 2014. Extensive behavioural divergence following colonisation of the freshwater environment in threespine sticklebacks. PLoS One 9: e98980.

Dingemanse, N.J., Plas F. Van Der, Wright, J., Réale, D., Schrama, M., Roff, D.A., et al. 2009. Individual experience and evolutionary history of predation affect expression of heritable variation in fish personality and morphology. Proc. R. Soc. B. 276: 1285–1293.

Dingemanse, N.J., Wright, J., Kazem, A.J.N., Thomas, D.K., Hickling, R. & Dawnay, N. 2007. Behavioural syndromes differ predictably between 12 populations of three-spined stickleback. J. Anim. Ecol. 76: 1128–1138.

Dingemanse, N., Dochtermann, N. & Wright, J. 2010. A method for exploring the structure of behavioural syndromes to allow formal comparison within and between data sets. Anim. Behav. 79: 439–450

Dingemanse, N., Barber, I. & Dochtermann, N. 2020. Non-consumptive effects of predation: does perceived risk strengthen the genetic integration of behaviour and morphology in stickleback? Ecol. Lett. 23: 107–118

Eriksson, B.K., Yanos, C., Bourlat, S.J., Donadi, S., Fontaine, M.C., Hansen, J.P., et al. 2021. Habitat segregation of plate phenotypes in a rapidly expanding population of three-spined stickleback. Ecosphere. 12(6): e03561.

Garcia-Elfring, A., Paccard, A., Thurman, T. J., Wasserman, B. A., Palkovacs, E. P., Hendry, A. P. et al. 2021. Using seasonal genomic changes to understand historical adaptation to new environments: Parallel selection on stickleback in highly-variable estuaries. Mol. Ecol. 30(9): 2054–2064.

Gingerich, P.D. 1993. Quantification and comparison of evolutionary rates. Am. J. Sci. 293A: 453–478.

Greenwood, A.K., Mills, M.G., Wark, A.R., Archambeault, S.L. & Peichel, C.L. 2016. Evolution of schooling behavior in threespine sticklebacks is shaped by the Eda gene. Genetics 203: 677–681.

Gross, M.R., Coleman, R.M. & McDowall, R.M. 1988. Aquatic productivity and the evolution of diadromous fish migration. Science. 239: 1291–1293.

Haldane, J.B.S. 1949. Suggestions as to quantitative measurement of rates of evolution. Evolution. 3: 51–56.

Hendry, A.P., Farrugia, T.J., Kinnison, M.T. 2008. Human influences on rates of phenotypic change in wild animal populations. Mol. Ecol. 17: 20–29.

Herczeg, G., Ab Ghani, N.I. & Merilä, J. 2013. Evolution of stickleback feeding behaviour: Genetics of population divergence at different ontogenetic stages. J. Evol. Biol. 26: 955–962.

Hoogland, R., Morris, D., & Tinbergen, N. 1956. The spines of sticklebacks (Gasterosteus and Pygosteus) as means of defence against predators (Perca and Esox). Behaviour. 10: 205–236.

Hosoki, T., Mori, S., Nishida, S., Kume, M., Sumi, T. & Kitano, J. 2019. Diversity of gill raker number and diets among stickleback populations in novel habitats created by the 2011 Tōhoku earthquake and tsunami. Evol. Ecol. Res. 20: 213–230.

Huizinga, M., Ghalambor, C.K. & Reznick, D.N. 2009. The genetic and environmental basis of adaptive differences in shoaling behaviour among populations of Trinidadian guppies, Poecilia reticulata. J. Evol. Biol. 22: 1860–1866.

Huntingford, F.A. & Wright, P.J. 1993. The development of adaptive variation in predator avoidance in freshwater fishes. Mar. Behav. Physiol. 23: 45–61.

Huntingford, F.A. 1976. The relationship between anti-predator behaviour and aggression among conspecifics in the three-spined stickleback, Gastrosteus aculeatus. Anim. Behav. 24: 245–260

Jones, F. C., Grabherr, M. G., Chan, Y. F., Russell, P., Mauceli, E., Johnson, J. et al. 2012. The genomic basis of adaptive evolution in threespine sticklebacks. Nature. 484: 55–61.

Kim, S. Y., & Velando, A. 2015. Phenotypic integration between antipredator behavior and camouflage pattern in juvenile sticklebacks. Evolution. 69(3): 830–838.

Kitano, J., Ishikawa, A., Kume, M. & Mori, S. 2012. Physiological and genetic basis for variation in migratory behavior in the three-spined stickleback, Gasterosteus aculeatus. Ichthyol. Res. 59: 293–303.

Kitano, J. & Lema, S.C. 2013. Divergence in thyroid hormone concentrations between juveniles of marine and stream ecotypes of the threespine stickleback (Gasterosteus aculeatus). Evol. Ecol. Res. 15: 143– 153.

Kitano, J., Lema, S.C., Luckenbach, J.A., Mori, S., Kawagishi, Y., Kusakabe, M., et al. 2010. Adaptive divergence in the thyroid hormone signaling pathway in the stickleback radiation. Curr. Biol. 20: 2124– 2130.

Kozak, G.M. & Boughman, J.W. 2012. Plastic responses to parents and predators lead to divergent shoaling behaviour in sticklebacks. J. Evol. Biol. 25: 759–769.

Kusakabe, M., Ishikawa, A., Ravinet, M., Yoshida, K., Makino, T., Toyoda, A., et al. 2017. Genetic basis for variation in salinity tolerance between stickleback ecotypes. Mol. Ecol. 26: 304–319.

Lacasse, J. & Aubin-Horth, N. 2014. Population-dependent conflict between individual sociability and aggressiveness. Anim. Behav. 87: 53–57.

Lam, T.J. & Hoar, W.S. 1967. Seasonal Effects of Prolactin on Freshwater Osmoregulation of the Marine Form (Trachurus) of the Stickleback (Gasterosteus Aculeatus). Can. J. Zool. 45: 509–516.

Lescak, E.A., Bassham, S.L., Catchen, J., Gelmond, O., Sherbick, M.L., von Hippel, F.A., et al. 2015. Evolution of stickleback in 50 years on earthquake-uplifted islands. PNAS. 112(52): E7204–E7212.

Marchinko, K.B. 2009. Predation’s role in repeated phenotypic and genetic divergence of armor in threespine stickleback. Evolution. 63: 127–138.

Marchinko, K.B. & Schluter, D. 2007. Parallel evolution by correlated response: Lateral plate reduction in threespine stickleback. Evolution. 61: 1084–1090.

Marchinko, K.B. & Schluter, D. 2007. Parallel evolution by correlated response: lateral plate reduction in threespine stickleback. Evolution. 61: 1084–1090.

McGhee, K.E., Feng, S., Leasure, S. & Bell, A.M. 2015. A female’s past experience with predators affects male courtship and the care her offspring will receive from their father. Proc. R. Soc. B 282: 20151840.

Moran, E.V., Hartig, F. & Bell, D.M. 2016. Intraspecific trait variation across scales: Implications for understanding global change responses. Glob. Chang. Biol. 22: 137–150.

Moran, N.P., Mossop, K.D., Thompson, R.M., Chapple, D.G. & Wong, B.B.M. 2017. Rapid divergence of animal personality and syndrome structure across an arid-aquatic habitat matrix. Oecologia 185: 55–67.

Réale, D., Reader, S.M., Sol, D., McDougall, P.T. & Dingemanse, N.J. 2007. Integrating animal temperament within ecology and evolution. Biol. Rev. 82: 291–318.

Reimchen, T. E., Bergstrom, C., & Nosil, P. 2013. Natural selection and the adaptive radiation of Haida Gwaii stickleback. Evol. Ecol. Res. 15: 241–269.

Reimchen, T.E. 1980. Spine deficiency and polymorphism in a population of Gasterosteus aculeatus:an adaptation to predators? Can. J. Zool. 58: 1232–1244.

Robinson, B.W. 2013. Evolution of growth by genetic accommodation in Icelandic freshwater stickleback. Proc. R. Soc. B Biol. Sci. 280. 20132197.

Sih, A. 2013. Understanding variation in behavioural responses to human-induced rapid environmental change: A conceptual overview. Anim. Behav. 85: 1077–1088.

Sih, A., Cote, J., Evans, M., Fogarty, S. & Pruitt, J. 2012. Ecological implications of behavioural syndromes. Ecol. Lett. 15: 278–289.

Sih, A., Ferrari, M.C.O. & Harris, D.J. 2011. Evolution and behavioural responses to human-induced rapid environmental change. Evol. Appl. 4: 367–387.

Snyder, R.J. 1991. Migration and life histories of the threespine stickleback: evidence for adaptive variation in growth rate between populations. Environ. Biol. Fishes 31: 381–388.

Sommer-Trembo, C., Petry, A.C., Gomes Silva, G., Vurusic, S.M., Gismann, J., Baier, J., et al. 2017. Predation risk and abiotic habitat parameters affect personality traits in extremophile populations of a neotropical fish (Poecilia vivipara). Ecol. Evol. 7: 6570–6581.

Stamps, J. & Groothuis, T.G.G. 2010. The development of animal personality: relevance, concepts and perspectives. Biol. Rev. 85: 301–325.

Stein, L.R. & Bell, A.M. 2014. Paternal programming in sticklebacks. Anim. Behav. 95: 165–171.

Stein, L.R. & Bell, A.M. 2019. The role of variation and plasticity in parental care during the adaptive radiation of three-spine sticklebacks. Evolution. 73: 1037–1044.

Tudorache, C., Blust, R. & De Boeck, G. 2007. Swimming capacity and energetics of migrating and non-migrating morphs of three-spined stickleback Gasterosteus aculeatus L. and their ecological implications. J. Fish Biol. 71: 1448–1456.

Ward, A.J.W., Thomas, P., Hart, P.J.B. & Krause, J. 2004. Correlates of boldness in three-spined sticklebacks (Gasterosteus aculeatus). Behav. Ecol. Sociobiol. 55: 561–568.

Wark, A.R., Wark, B.J., Lageson, T.J. & Peichel, C.L. 2011. Novel methods for discriminating behavioral differences between stickleback individuals and populations in a laboratory shoaling assay. Behav. Ecol. Sociobiol. 65: 1147–1157.

Wolf, M. & Weissing, F.J. 2012. Animal personalities: consequences for ecology and evolution. Trends Ecol. Evol. 27: 452–461.

Wolf, M., Van Doorn, G.S. & Weissing, F.J. 2011. On the coevolution of social responsiveness and behavioural consistency. Proc. R. Soc. B Biol. Sci. 278: 440–448. Royal Society.

Wolf, M., Van Doorn, G.S., Weissing, F.J., Doorn G.S. van & Weissing, F.J. 2008. Evolutionary emergence of responsive and unresponsive personalities. Proc. Natl. Acad. Sci. 105: 15825–15830.

Wund, M.A., Singh, O.D., Geiselman, A. & Bell, M.A. 2012. Morphological evolution of an anadromous threespine stickleback population within one generation after reintroduction to Cheney Lake, Alaska. Evol. Ecol. Research. 17: 203–224

WWF (2020) Living Planet Report 2020 -Bending the curve of biodiversity loss. Almond, R.E.A., Grooten M. and Petersen, T. (Eds). WWF, Gland, Switzerland.

